# GLH/VASA helicases promote germ granule formation to ensure the fidelity of piRNA-mediated transcriptome surveillance

**DOI:** 10.1101/2022.01.21.477267

**Authors:** Wenjun Chen, Jordan S Brown, Tao He, Wei-Sheng Wu, Shikui Tu, Zhiping Weng, Donglei Zhang, Heng-Chi Lee

## Abstract

The ability to distinguish non-self from self is the key characteristic for any defense system. piRNAs function as guardians of the genome by silencing non-self nucleic acids and transposable elements in animals. Many piRNA factors are enriched in perinuclear germ granules, but whether their localization is required for piRNA biogenesis or function is not known. Here we show that GLH/VASA helicase mutants exhibit defects in forming perinuclear condensates containing PIWI and other small RNA cofactors. These mutant animals produce largely normal levels of piRNA but are defective in triggering piRNA silencing. Strikingly, while many piRNA targets are activated in GLH mutants, we observed that hundreds of endogenous genes are aberrantly silenced by piRNAs. This defect in self versus non-self recognition was also observed in other mutants where perinuclear P granules are disrupted. Together, our results argue that perinuclear germ granules function critically to promote the fidelity of piRNA-based transcriptome surveillance in *C. elegans* and preserve self versus non-self distinction.

## INTRODUCTION

Argonaute proteins use their associated small RNAs as guides to regulate targets with complementary sequences (Hutvagner and Simard 2008). The PIWI Argonaute and its associated piRNAs are conserved guardians of the animal genome that repress transposons in the germline (Batista et al. 2008; Brennecke et al. 2007; Carmell et al. 2007; Das et al. 2008; Saito 2006). A prerequisite for any defense system is the ability to distinguish non-self from self. In *C. elegans,* piRNAs trigger gene silencing of non-self RNAs through the recruitment of RNA-dependent RNA Polymerases (RdRPs) to produce WAGO Argonaute-associated 22G-RNAs that mediate transcriptional and posttranscriptional gene silencing (Gu et al. 2009; Shirayama et al. 2012). While diverse PIWI/piRNAs can recognize both foreign nucleic acids and germline-expressed “self” mRNAs (Shen et al. 2018)(Zhang et al. 2018), self RNAs are protected from piRNA silencing by Argonaute CSR-1 and its associated 22G-RNAs (Wedeles et al. 2013)(Seth et al. 2013) (Shen et al. 2018).

Intriguingly, the PIWI-related PRG-1, WAGO-1 and CSR-1 Argonautes are all enriched in germ granules, also known as P granules in *C. elegans* (Strome and Wood 1982)(Batista et al. 2008; Claycomb et al. 2009; Gu et al. 2009). Germ (P) granules are phase-separated liquid droplets that are found in the germ cells of all animals (Eddy 1975)((Brangwynne et al. 2009). The localization and formation of P granules are tightly controlled during *C. elegans* development (Updike and Strome 2010)(Seydoux 2018). In early embryos, P granules are cytoplasmic and sort to daughter cells of the germ cell lineage. As zygotic transcription begins later in embryogenesis, P granules re-localize to the nuclear periphery and remain perinuclear throughout most of germline development.

Mutations that affect the formation of cytoplasmic P granules in embryos, such as mutants for germ plasm factors *meg-3 meg-4*, impact the potency and inheritance of RNAi. (Lev et al. 2019)(Dodson and Kennedy 2019)(Ouyang et al. 2019). However, *meg-3 meg-4* mutant adults have normal perinuclear P granules. The role perinuclear P granules play in small RNA-mediated gene regulation is less clear. Several lines of evidence suggest that perinuclear P granules may be the site of mRNA surveillance by small RNAs. First, perinuclear P granules have been shown to be the major sites of mRNA transport in germline nuclei (Sheth et al. 2010). Second, the size of perinuclear P granules shrink soon after inhibition of mRNA transcription or mRNA export (Sheth et al. 2010)(Chen et al. 2020), consistent with the model newly exported mRNAs gather in perinuclear P granules. Third, a recent report showed that tethering an mRNA to P granule component PGL-1 leads to its silencing (Aoki et al. 2021), suggesting the accumulation of mRNA in P granules can trigger silencing by small RNA pathways. In addition, the enzymes required for piRNA 3’ end trimming (PARN-1) and the biogenesis of CSR-1 and WAGO associated 22G-RNAs (EGO-1) are both enriched in perinuclear P granules (Tang et al. 2016)(Phillips et al. 2012). Together, these observations raise the possibility that the production and/or function of small RNAs may require the enrichment of these factors in perinuclear P granules.

The VASA-homologue RNA helicases GLH-1 and GLH-4 play a critical role in the formation of both cytoplasmic and perinuclear P granules. (Roussell and Bennett 1993)(Gruidl et al. 1996)(C. Spike et al. 2008)(Elisabeth A. Marnik, J. Heath Fuqua, Catherine S. Sharp, Jesse D. Rochester, Emily L. Xu, Sarah E. Holbrook 2019)(Chen et al. 2020). In addition, several other P granule factors, including DEPS-1 and PGL-1, have been reported to promote both cytoplasmic and perinuclear P granule assembly (Hanazawa et al. 2011; C. A. Spike et al. 2008). These observations make them possible candidates as arbiters for examining P granule function in small RNA mediated gene silencing.

Here we show that GLH-1 and GLH-4 play a global role in promoting the liquid condensation of Argonautes and other small RNA factors at perinuclear foci. In addition, we found that the biogenesis of neither piRNAs nor secondary small RNAs, including WAGO or CSR-1 associated 22G-RNAs, broadly require GLH/VASA. In GLH and in other mutants with defects in forming perinuclear P granules, many piRNA targets were activated, with fewer secondary WAGO 22G- RNAs produced at piRNA targeting sites. Additionally, many functional endogenous mRNAs are aberrantly silenced by piRNAs. Together, our results suggest that GLH/VASA helicases and perinuclear P granules are critical for ensuring the fidelity of mRNA surveillance by piRNAs and that without P granules, small RNA pathways can no longer robustly identify mRNAs as self or non-self.

## RESULTS

### GLH/VASA helicases promote the liquid condensation of piRNA pathway components throughout germline development

While GLH-1 plays a critical role in controlling the perinuclear localization of PIWI PRG-1 (Elisabeth A. Marnik, J. Heath Fuqua, Catherine S. Sharp, Jesse D. Rochester, Emily L. Xu, Sarah E. Holbrook 2019)(Chen et al. 2020), we failed to detect PRG-1 in GLH-1 complexes by mass spectrometry under native conditions (Chen et al. 2020). We hypothesized that their interaction could be transient and therefore applied the chemical crosslinking reagent dithio-bis- maleimidoethane (DTME) to capture potentially transient interactions (Ge et al. 2019). Indeed, using DTME-crosslinked worms, we are able to detect PRG-1 in the GLH-1 complex (Figure 1A). In addition, we observed many other small RNA components in the GLH-1 complex, including P granule factors DEPS-1, WAGO- 1, CSR-1, and Z granule factor WAGO-4 (Figure 1A and Table S1). Z granules are derived from P granules during embryogenesis and remain adjacent to P granules in the adult germline (Wan et al. 2018)(Ishidate et al. 2018). These observations raise the possibility that GLH helicases may play a global role in regulating the localization of small RNA pathway components. As VASA-like helicases GLH-1 and GLH-4 function redundantly to promote the localization of PIWI PRG-1 (Chen et al. 2020), we examined the localization of various small RNA machinery in the *glh-1 glh-4* mutant. Consistent with previous findings, we found that for PIWI PRG-1, both perinuclear and cytoplasmic foci are greatly reduced in the *glh-1 glh-4* double null mutant (Figure 1B and S1A). For several other small RNA factors, including P granule factors - DEPS-1, CSR-1, and EGO-1, and Z granule factors - WAGO-4 and ZFNX-1, their perinuclear localization is also significantly reduced in the *glh-1 glh-4* double mutant, although residual foci of some of these factors can still be observed (Figure 1B, C and S1A, B). We then examined the localization of MUT-16, a key factor in the assembly of Mutator granules (Phillips et al. 2012). Mutator granules house the small RNA components involved in producing WAGO associated 22G-RNAs.

**Figure 1.**
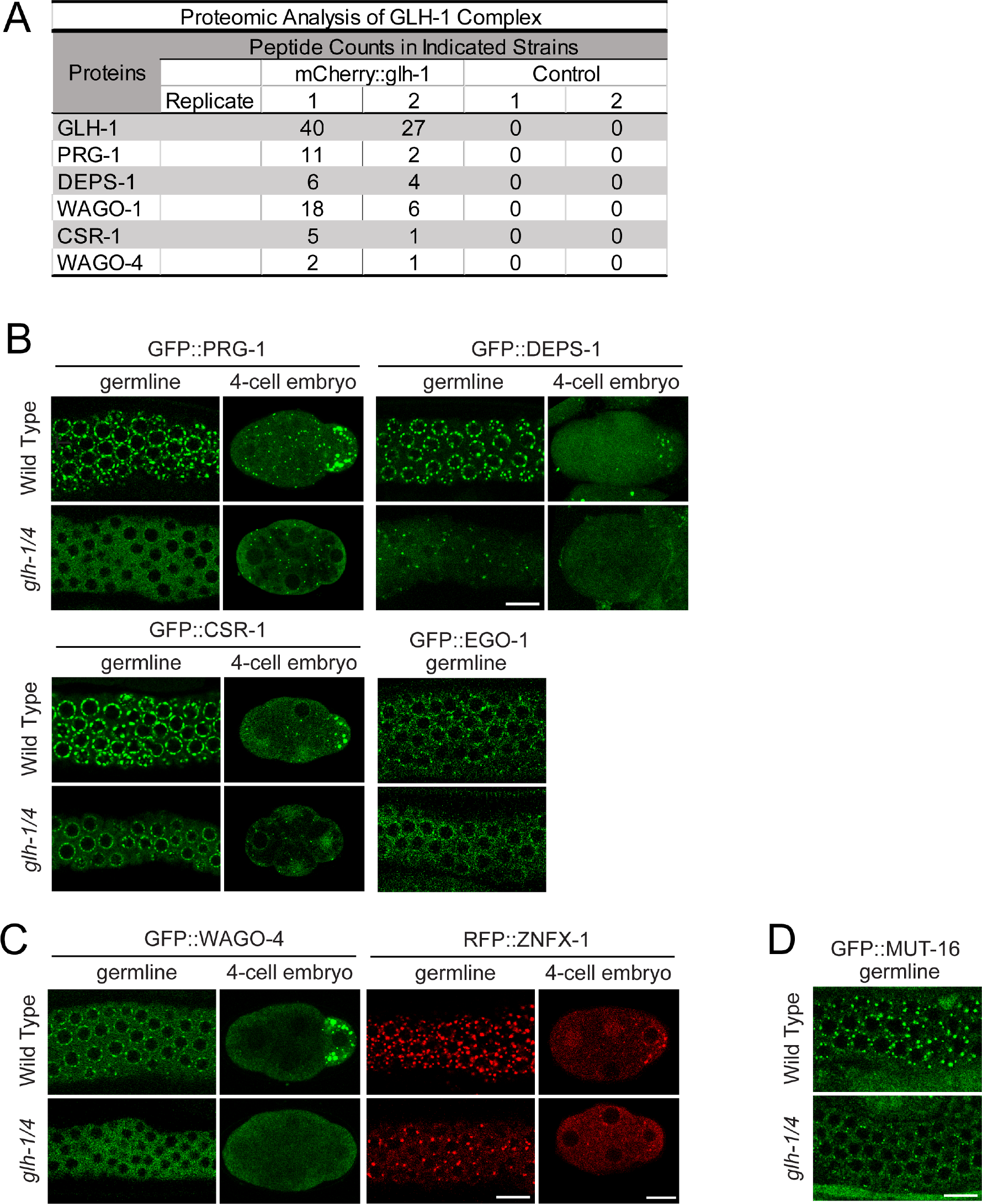
The GLH/VASA helicases GLH-1 and GLH-4 promote the localization of germ granule factors. (a) Proteomic analyses of GLH-1 complex. The numbers of peptides identified in two independent pull-down experiments are shown. (b) Fluorescent micrographs show the localization of the specific proteins known to be enriched in P granules in the wild type or the *glh-1 glh-4* mutant in the adult germline (left) and in the four-cell embryo (right). Bar, 10 micrometers. (c) Fluorescent micrographs show the localization of the specific proteins known to be enriched in Z granules in the wild type or the *glh-1 glh-4* mutant in the adult germline (left) and in the four-cell embryo (right). Bar, 10 micrometers. (d) Fluorescent micrographs show the localization of the specific proteins known to be enriched in mutator granules in the wild type or the *glh-1 glh- 4* mutant in the adult germline. Bar, 10 micrometers.

Mutator granules are frequently found in close contact with perinuclear P granules (Celja J. Uebel, Dana Agbede, Dylan C. Wallis 2020). Previous experiments using RNAi knockdown of *glh-1* and *glh-4* did not lead to disruption of MUT-16 localization (Phillips et al. 2012). Surprisingly, the localization of MUT- 16 was significantly disrupted in *glh-1 glh-4* double mutants (Figure 1D and S1C). Together, these results show that GLH/VASA helicases play a global role in enriching small RNA machinery into the distinct liquid condensates observed throughout *C. elegans* germline development, including cytoplasmic and perinuclear P granules, Z granules and Mutator granules.

### GLH/VASA helicase mutants exhibit defects in piRNA silencing

To investigate whether GLH-1 and GLH-4 helicases are required for piRNA-mediated gene silencing, we examined whether the silencing of a piRNA reporter (Seth et al. 2018) requires these GLH/VASA helicases. This piRNA reporter is silenced in wild type animals but is activated in the *prg-1* mutant background (Figure 2A). Similarly, we found that the piRNA reporter is activated in the *glh-1 glh-4* double mutant background (Figure 2A), suggesting GLH-1 and GLH-4 play a role in piRNA silencing. In addition, we found that the piRNA reporter is also activated in the *glh-1 DQAD* mutant background. In the *glh-1 DQAD* mutant, large PRG-1 and WAGO-4 aggregates are found in the cytoplasm with a significant reduction in perinuclear PRG-1 and WAGO-4 foci (Figure 2B). In addition, these abnormal, cytoplasmic aggregates are not properly sorted to the germ cell lineage, leading to the presence of these foci in somatic lineages (Figure 2B). Together, these data show that GLH mutants are defective in piRNA silencing.

**Figure 2.**
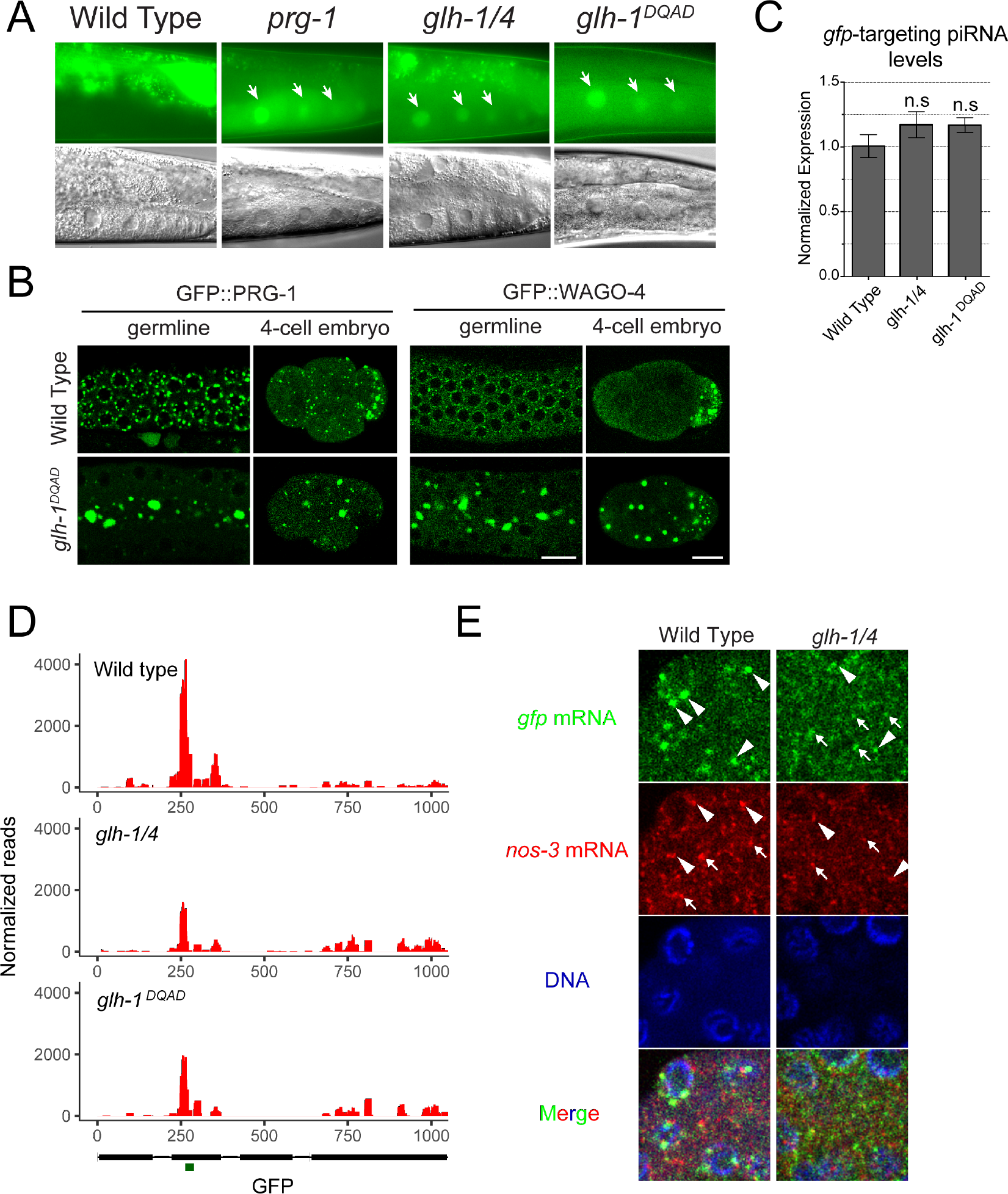
GLH/VASA helicases are required for piRNA-dependent silencing of a piRNA reporter. (A) GFP expression of the piRNA reporter in the indicated strains. Arrows indicate the adult germline nuclei in maturing oocytes with expression of GFP transgenes (top). DIC images of the corresponding reporter images (bottom). (B) Fluorescent micrographs show the localization of PRG-1 and WAGO-4 in the indicated strains in the adult germline (left) and in the four-cell embryo (right). Bar, 10 micrometers. (C) q-RT PCR measurements of the abundance of the GFP-targeting piRNA in the indicated strains. Statistical analysis was performed using a two- tailed Student’s t-test. (D) 22G-RNAs distribution at GFP coding sequences in the indicated strains. The green bar (below) indicates the location of the GFP sequence complementary to the GFP-targeting piRNA. (E) Photomicrographs of pachytene nuclei in fixed adult gonads hybridized with single molecule fluorescent (smFISH) probes complementary to *gfp* mRNA (Green) and *nos-3* mRNA (Red) in the indicated strains. Nuclear DNA was stained with DAPI (Blue). The arrowheads indicate perinuclear mRNA foci while arrows indicate cytoplasmic mRNA foci.

We then wanted to understand why piRNA silencing is defective in GLH/VASA mutants. We found no significant change in expression of the GFP-targeting piRNA in these GLH/VASA mutants compared to wild type animals (Figure 2C). In contrast, we observed a pronounced reduction in the 22G-RNAs produced around the GFP-targeting piRNA binding site in both the *glh-1 glh-4* double and *glh-1 DQAD* mutants (Figure 2D). These analyses suggest that GLH- 1 and GLH-4 are not required for the biogenesis of the GFP-targeting piRNA, but rather promote the production of 22G-RNAs at the piRNA targeting site.

Since perinuclear P granules have been shown to be the major sites of mRNA export (Sheth et al. 2010), we wondered whether GLH/VASA may also contribute to the localization of target mRNAs. We therefore examined the localization of GFP mRNAs, the piRNA target of our piRNA reporter, using single molecule fluorescent *in situ* hybridization (smFISH). In our piRNA reporter strain where GFP is silenced by a GFP-targeting piRNA, we observed large GFP perinuclear foci and little cytoplasmic GFP signal (Figure 2E). In the *glh-1 glh-4* double mutant piRNA reporter strain where GFP is activated, the perinuclear GFP foci were greatly reduced, while more cytoplasmic signal was observed (Figure 2E). Therefore, GLH-1 and GLH-4 promotes the location of GFP mRNA at perinuclear foci in the piRNA reporter. These observations are consistent with the model that GLH/VASA promotes the accumulation of PRG-1, piRNA cofactors, and target RNAs at perinuclear foci to trigger piRNA silencing.

### Mutants defective in P granule assembly exhibit defects in piRNA silencing

The GLH mutants examined above are defective in the localization of small RNA machinery in both perinuclear and cytoplasmic P granules. To examine whether piRNA silencing capability correlates with the ability to form either type of liquid condensate, we characterized additional genetic mutants that have been reported to show defects in forming either cytoplasmic and/or perinuclear P granules. The P granule component DEPS-1 has been shown to promote the assembly of perinuclear and cytoplasmic P granules (C. A. Spike et al. 2008).

Indeed, the formation of both perinuclear and cytoplasmic WAGO-4 condensates are compromised in *deps-1* mutants (Figure 3A). However, we found *deps-1* mutants exhibit a reduced number of perinuclear PRG-1 condensates but retained normal cytoplasmic PRG-1 condensates. While we have not yet confirmed this finding using our piRNA reporter, a recent study reports that DEPS-1 is required for silencing of a piRNA reporter (Suen et al. 2020). In addition, we have recently shown that the N terminal phenylalanine-glycine- glycine (FGG) repeats of GLH-1 promote its perinuclear localization, leading to the recruitment of PIWI PRG-1 at perinuclear P granules (Chen et al. 2020).

**Figure 3.**
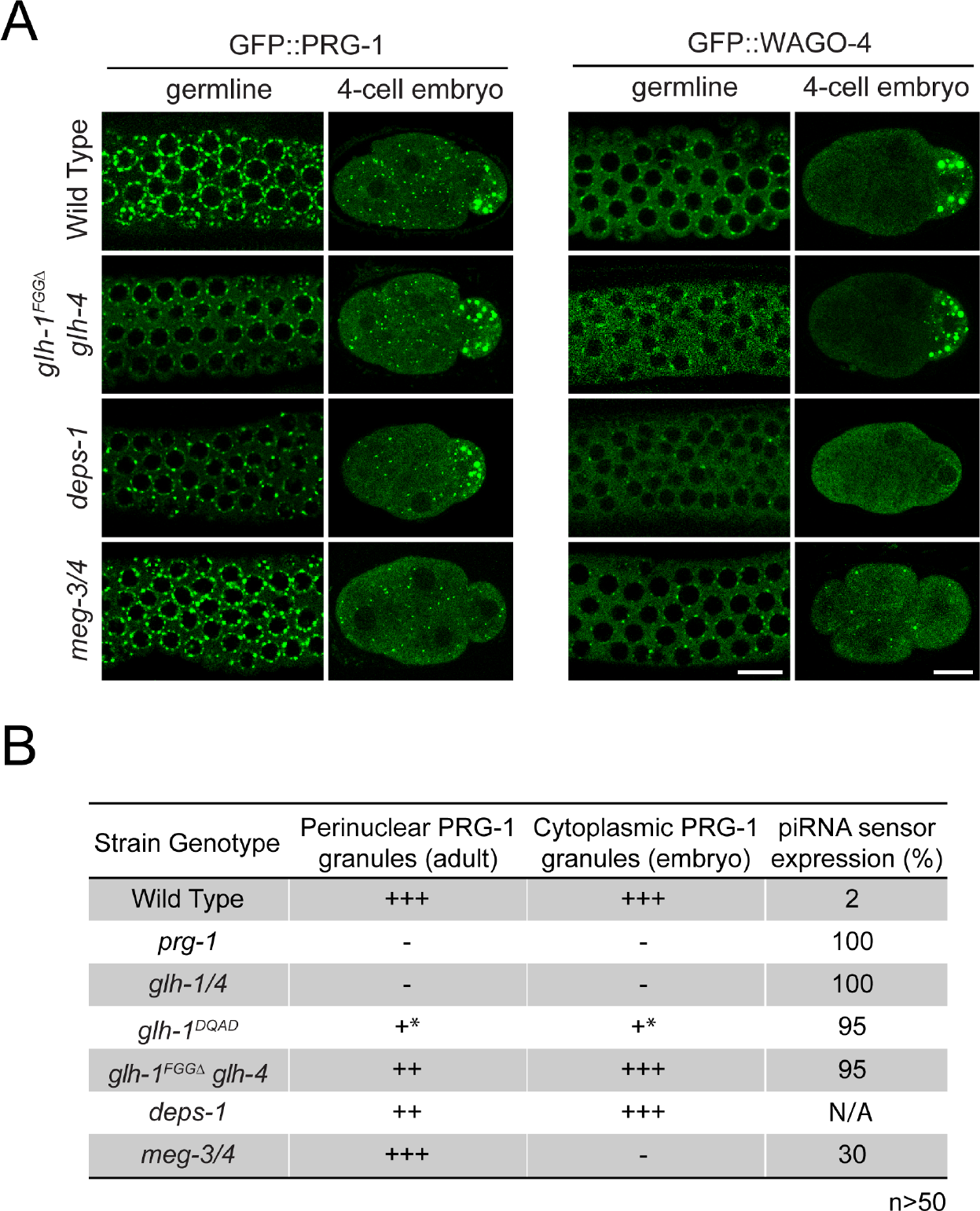
Mutants with defective in either perinuclear or cytoplasmic P granules exhibit defects in gene silencing. (A) Fluorescent micrographs show the localization of PRG-1 and WAGO-4 in the indicated strains in the adult germline (left) and in the four-cell embryo (right). Bar, 10 micrometers. (B) GFP expression in the piRNA reporter and the ability to form perinuclear or cytoplasmic PRG-1 granules in the indicated strains.

Indeed, we confirmed that PRG-1 and WAGO-4 perinuclear condensates, but not their cytoplasmic condensates, are partially disrupted in the *glh-1* FGGΔ *glh-4* mutant (Figure 3A). We also found that the piRNA reporter is activated in this *glh- 1 FGGΔ glh-4* strain (Figure 3B). These results indicate that mutants defective in forming perinuclear P granules exhibit defects in piRNA silencing. We then examined the *meg-3 meg-4* mutant, in which the localization of the small RNA machinery is only disrupted in cytoplasmic P granules (Smith et al. 2016)(Ouyang et al. 2019) (Figure 3A). We found that the piRNA reporter is also activated in ∼30 percent of *meg-3 meg-4* mutant animals (Figure 3B). Together, these results indicate that the localization of piRNA factors at perinuclear and cytoplasmic P granules can both contribute to their function in piRNA silencing.

### Mutants with abnormal perinuclear P granules exhibit defects in the initiation of piRNA silencing

Our observations that GLH/VASA promotes the perinuclear localization of piRNA factors and their target mRNAs raises the possibility that their enrichment in P granules allows PIWI PRG-1 and its cofactors to efficiently identify their targets and to trigger gene silencing. To test whether GLH/VASA mutants exhibit defects in *de novo* piRNA-mediated gene silencing, we microinjected a synthetic piRNA-expressing plasmid into transgenic worms expressing a silencing-prone GFP::CDK-1 transgene (Zhang et al 2018) and monitored the silencing of the GFP transgene by the GFP-targeting piRNA (Figure 4A). While 97% of the injected wild type animals successfully triggered silencing of the GFP transgene by the GFP-targeting piRNA by the F2 generation, only 5% of the *glh-1 glh-4* double mutant animals and 36% of the *glh-1 DQAD* mutant animals triggered silencing of the GFP transgene (Figure 4B). In addition, we observed that only 40% of the *deps-1* mutant animals and 31% of the *glh-1 FGGΔ glh-4* mutant animals triggered silencing of the GFP transgene. In contrast, *meg-3 meg-4* mutants exhibit a normal, wild type ability to trigger silencing of the GFP transgene by the GFP-targeting piRNA. Taken together, these assays show that mutants defective in forming perinuclear P granules are compromised in their ability to initiate piRNA silencing.

**Figure 4.**
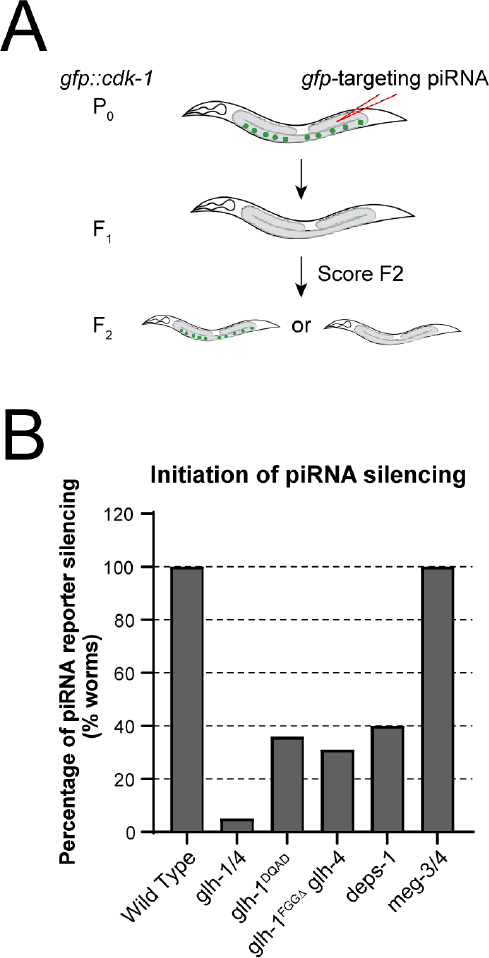
Mutants with defects in PRG-1 perinuclear condensates exhibit defects in triggering piRNA silencing. (A) A schematic showing the assay for examining the *de novo* silencing of a GFP transgene by a GFP-targeting piRNA. (B) The percentage of F2 generation worms that exhibit silencing of a GFP transgene in the indicated strains.

### Mutants defective in perinuclear P granules exhibit defects in silencing endogenous piRNA targets

Our analyses of the piRNA reporter suggest that GLH/VASA is not required for the biogenesis of a GFP-targeting piRNA (Figure 2C). Consistent with the reporter analyses, we observed slightly reduced or normal piRNA levels in the *glh-1 glh-4* double mutant or in *glh-1 DQAD* mutant, respectively (Figure S3A). In addition, we found that the length of piRNAs was not changed in the *glh-1 glh-4* double mutant or in the *glh-1 DQAD* mutant (Figure S3B), suggesting that the 3’ processing of piRNAs by PARN-1 is not affected. On the other hand, over 41% of WAGO targets (genes known to be silenced by WAGO 22G-RNAs) exhibit a two-fold decrease in 22G-RNAs in the *glh-1 glh-4* double mutant and over 51% in the *glh-1 DQAD* mutant (Figure 5A, left and Figure S3C) (Gu et al. 2009). To confirm that the reduction in 22G-RNAs is due to decreased production of WAGO-associated 22G-RNAs, we performed an immunoprecipitation with HRDE-1 (WAGO-9) and compared wild type HRDE-1 22G-RNAs to those in *glh-1* mutants, which exhibit a partial defect in forming perinuclear PRG-1 foci (Chen et al. 2020). (Figure 5A, right). To obtain a more conservative list of affected WAGO genes, we also applied a statistical threshold and observed that 16% of WAGO genes were significantly reduced in HRDE-1 22G-RNA accumulation in *glh-1* mutants, compared to only 5% of WAGO genes significantly increased. Using mRNA sequencing, we found the majority of genes exhibiting significantly reduced HRDE-1 22G-RNA levels in *glh-1* mutants had increased mRNA levels in *glh-1 glh-4* double mutants and in the *glh-1 DQAD* mutants (Figure 5B and Figure S3D). Several transposons were among those shared activated genes, including TC2 and PAL8C_1 (Figure S3E). These observations indicate that GLH/VASA mutants exhibit defects in silencing WAGO targets. As the levels of 22G RNAs from WAGO targets were reported to be reduced in *meg-3 meg-4* double mutants (Ouyang et al. 2019) which exhibit defects only in cytoplasmic but not in perinuclear PRG-1 granules, we wondered whether distinct or common WAGO targets exhibit 22G-RNA defects in mutants defective in cytoplasmic versus perinuclear P granules. We noticed that those WAGO genes with significantly reduced HRDE-1 22G-RNAs in *glh-1* mutants exhibit a greater reduction in 22G-RNAs and a greater increase in mRNA expression compared to other WAGO target genes in the *glh-1 glh-4* mutants and in *glh-1* DQAD mutants.

**Figure 5.**
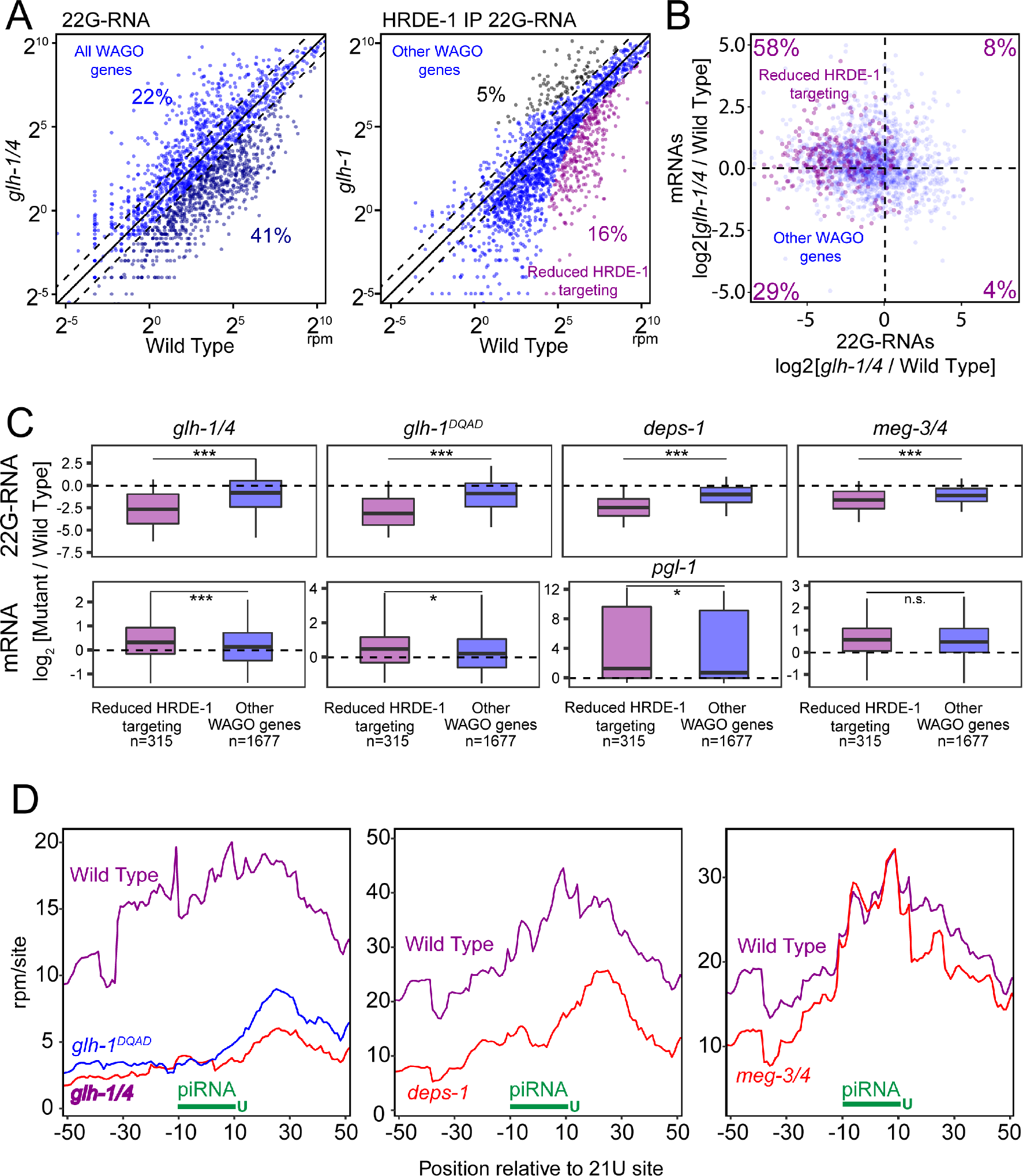
GLH/VASA mutants exhibit defects in producing secondary WAGO-22G-RNAs. (A) A scatter plot showing the abundance of all 22G-RNAs (left) or HRDE-1 bound 22G-RNAs (right) mapped to each WAGO target in wild type worms compared to indicated strains. (Left) The percentage of WAGO targets with 2-fold increased or decreased 22G-RNAs in mutants are shown. (Right) The percentage of significantly changed (*P* < 0.05 and 2- fold) WAGO targets are shown. The three diagonal lines indicate a two- fold increase (top), no change (middle), or a two-fold depletion (bottom) in the indicated mutant strains. (B) A scatter plot showing the mRNA vs 22G-RNA log2 expression changes for WAGO targets in *glh-1 glh-4* mutants vs wild type worms. The upper left quadrant corresponds to WAGO targets that have become activated in the mutant (increased mRNAs and decreased 22G-RNAs). Percentages are shown in each quadrant to indicate the proportion of WAGO targets with reduced HRDE-1-associated 22G-RNAs in the *glh-1* mutant that fall in that quadrant. (C) 22G-RNA and mRNA fold changes for WAGO targets with reduced HRDE-1-associated 22G-RNAs in the *glh-1* mutant versus all other WAGO targets in the indicated strains versus wild type worms. Statistical analysis was performed using a two-tailed Mann-Whitney Wilcoxon test. *** indicates *P* < 0.001; * indicates *P* < 0.05. (D) Density of 22G-RNAs within a 100 nt window around predicted piRNA target sites in the indicated strains. Computed by summing 22G-RNA density per piRNA targeting site in WAGO targets with reduced HRDE-1 associated 22G-RNAs in the *glh-1* mutant. The plots are centered on the 10^th^ nucleotide of piRNAs.

A similar trend of 22G-RNA changes is also found in *deps-1* mutants. In addition, in the mutant for PGL-1, another P granule factor that promotes the localization of GLH-1 and WAGO-1 (Aoki et al. 2021), mRNA expression for those reduced HRDE-1 22G-RNA target genes found in *glh-1* mutants also increased more than other WAGO genes (Knutson et al 2017). (Figure 5C). In *meg-3 meg-4* mutants, which exhibit defects in the formation of cytoplasmic P granules, we see much less difference and no difference in 22G-RNA and mRNA levels between these WAGO genes, respectively (Figure 4C). These results suggest that hundreds WAGO target genes are preferentially activated in GLH/VASA mutants and other mutants exhibiting defects in perinuclear P granules.

Since our piRNA reporter analyses suggest that GLH/VASA is critical for the production of downstream 22G-RNAs around piRNA targeting sites, we wondered whether the production of piRNA-dependent 22G-RNAs are preferentially affected in GLH/VASA mutants and other mutants exhibiting defects in forming perinuclear P granules (Lee et al. 2012). Indeed, in *glh-1 glh-4, glh-1* DQAD and *deps-1* mutants, those WAGO target genes in which the production of 22G-RNAs most depends on PRG-1 exhibit greater reductions in 22G-RNAs than other WAGO targets (Figure S3F). A still significant but lesser difference in 22G-RNA production was found for *meg-3 meg-4* mutants (Figure S3F). To specifically examine the production of 22G-RNA production at piRNA targeting sites, we compared the local production of 22G-RNAs at predicted piRNA target sites (Wu et al. 2019; Zhang et al. 2018). For those WAGO targets with significantly reduced HRDE-1 22G-RNAs in *glh-1* mutants, 22G-RNAs are enriched around predicted piRNA target sites in wild type but much less so in *glh- 1 glh-4*, *glh-1 DQAD*, or *deps-1* mutants (Figure 5D). In contrast, while 22G- RNAs are overall slightly reduced in *meg-3 meg-4* mutants, 22G-RNAs remain enriched around these predicted piRNA target sites (Figure 4E). Similar results were observed when we looked at predicted piRNA target sites in all WAGO genes (Figure S3G). These observations suggest that the production of piRNA-dependent 22G-RNAs is preferentially compromised in the *glh* and *deps-1* mutants. Together, these observations suggest that piRNA-dependent WAGO target genes are preferentially activated in VASA mutants and other mutants exhibiting defects in forming perinuclear P granules.

### Mutants defective in the formation of perinuclear P granules aberrantly silence many functional endogenous genes

Essentially all germline transcripts are targeted either by 22G-RNAs associated with WAGO or CSR-1 Argonautes, which can silence or license the expression of their targets, respectively (Gu et al. 2009)(Wedeles et al. 2013)(Seth et al. 2013). Since CSR-1 is present in GLH-1 complexes and the formation of CSR-1 perinuclear condensates is promoted by GLHs (Figure 1A&B), we wondered whether GLHs may regulate the biogenesis and/or function of CSR-1-associated 22G-RNAs. 22G-RNAs antisense to CSR-1 target genes remain mostly unchanged in *glh* mutants compared to wild type (Figure 6A, left and Figure S4A). Intriguingly, in *glh-1 glh-4* double mutants, we noticed that 717 CSR-1 genes (15%) exhibit at least a two-fold increase in 22G-RNAs that mapped to them (Table S2). Strikingly, HRDE-1 (WAGO-9) immunoprecipitation analyses of the *glh-1* mutant showed that 320 genes were significantly enhanced in HRDE-1 22G-RNA accumulation in *glh-1* mutants. mRNA sequencing analyses confirmed an overall reduction in mRNA levels for these genes in *glh-1 glh-4* double and in *glh-1 DQAD* mutants (Figure 6B and Figure S4A). The aberrantly silenced CSR-1 targets include several functional genes involved in germline development, including *nos-2*, *puf-6* and *puf-7* (Table S2).

**Figure 6.**
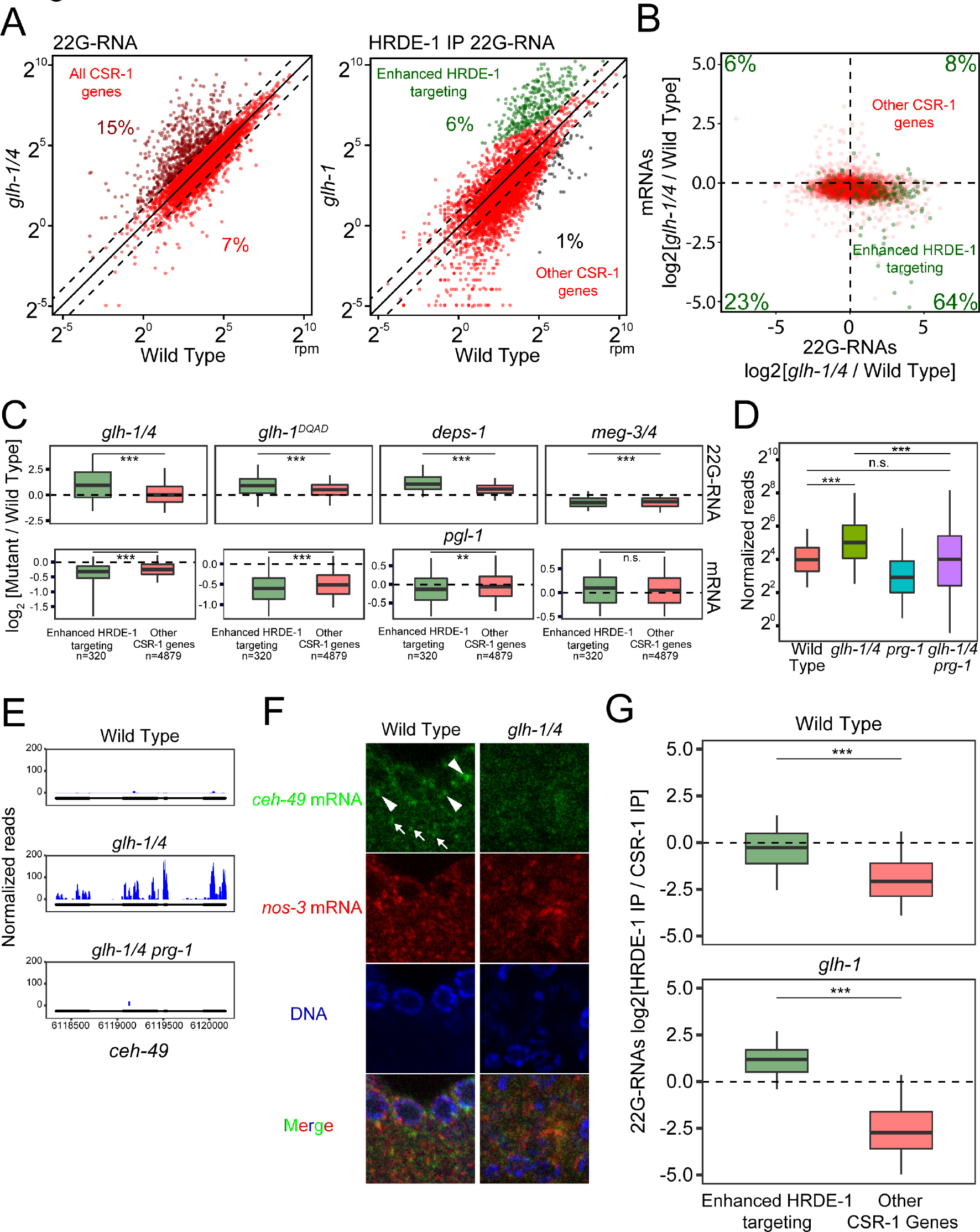
Many functional germline genes are silenced in mutants with defects in forming perinuclear PRG-1 granules. (A) A scatter plot showing the abundance of all 22G-RNAs (left) or HRDE-1 bound 22G-RNAs (right) mapped to each CSR-1 target in wild type worms compared to indicated strains. (Left) The percentage of CSR-1 targets with 2-fold increased or decreased 22G-RNAs in mutants are shown. (Right) The percentage of significantly changed (*P* < 0.05 and 2-fold) CSR-1 targets are shown. The three diagonal lines indicate a two-fold increase (top), no change (middle), or a two-fold depletion (bottom) in the indicated mutant strains. (B) A scatter plot showing the mRNA vs 22G-RNA log2 expression changes for CSR-1 targets in *glh-1 glh-4* mutants vs wild type worms. The bottom right quadrant corresponds to CSR-1 targets that have become silenced in the mutant (decreased mRNAs and increased 22G-RNAs). Percentages are shown in each quadrant to indicate the proportion of CSR-1 targets with enhanced HRDE-1-associated 22G-RNAs in the *glh-1* mutant that fall in that quadrant. (C) 22G-RNA and mRNA fold changes for CSR-1 targets with enhanced HRDE-1-associated 22G-RNAs in the *glh-1* mutant versus all other CSR-1 targets in the indicated strains versus wild type worms. Statistical analysis was performed using a two-tailed Mann-Whitney Wilcoxon test. *** indicates *P* < 0.001; ** indicates *P* < 0.01. (D) 22G-RNA accumulation for CSR-1 targets with enhanced HRDE-1- associated 22G-RNAs in the *glh-1* mutant compared between the indicated strains. Statistical analysis was performed using a two-tailed Mann-Whitney Wilcoxon test. *** indicates *P* < 0.001. (E) 22G-RNAs distribution in the enhanced HRDE-1 associated 22G-RNA target *ceh-49* in the indicated strains. (F) Photomicrographs of pachytene nuclei in fixed adult gonads hybridized with single molecule fluorescent (smFISH) probes complementary to *ceh-49* mRNA (Green) and *nos-3* mRNA (Red) in the indicated strains. Nuclear DNA was stained with DAPI (Blue). The arrowheads indicate perinuclear mRNA foci while arrows indicate cytoplasmic mRNA foci. (G) 22G-RNA enrichment in HRDE-1 versus CSR-1 IP experiments in the indicated strains for CSR-1 targets and WAGO targets. The dotted line indicated no enrichment for either Argonaute. Log2 enrichment above 0 indicates HRDE-1 dominance while below 0 indicates CSR-1 dominance. Statistical analysis was performed using a two-tailed Mann-Whitney Wilcoxon test. *** indicates *P* < 0.001.

We wondered whether the aberrant silencing of CSR-1 genes occurs in other mutants defecting in cytoplasmic and/or perinuclear P granule formation. In *glh-1 DQAD* mutants, a significant increase in 22G-RNAs and decrease in mRNAs are found for those CSR-1 genes with enhanced HRDE-1 22G-RNA levels in *glh-1* mutants. Similar changes of 22G-RNA or mRNA levels are found in *deps-1* mutant or *pgl-1* mutant, respectively. In contrast, in the *meg-3 meg-4* mutant that only disrupts the formation of cytoplasmic P granules in embryos, we did not observe an increase in 22G-RNAs nor a decrease in mRNAs for these CSR-1 genes with enhanced HRDE-1 22G-RNA levels in *glh-1* mutants (Figure 6C). These observations suggest that in mutants with abnormal perinuclear P granules, a group of CSR-1 target genes are aberrantly silenced by WAGO 22G- RNAs.

### PRG-1 is required for the aberrant CSR-1 gene silencing in VASA mutants

Because perinuclear P granule loss affects the localization but not abundance of PIWI PRG-1 (Chen et al. 2020), we wondered whether PRG-1 may be aberrantly recognizing these CSR-1 genes when perinuclear P granule formation is compromised. Indeed, we found that the aberrant production of 22G- RNAs against CSR-1 genes with enhanced HRDE-1 22G-RNA levels in *glh-1* mutants are significantly reduced in *glh-1 glh-4 prg-1* triple mutants compared to *glh-1 glh-4* double mutants (Figure 6D). The mutation of *prg-1* suppresses the aberrant accumulation of CSR-1 22G-RNAs in *glh-1 glh-4* double mutants by 2- fold or more for nearly two-thirds of affected genes (Figure S4B). These data suggest that most of the aberrant mRNA silencing of CSR-1 genes found in GLH/VASA mutants is triggered by piRNAs. Together, these data suggest VASA not only promotes silencing of non-self (WAGO targets), but also promotes licensing of self (CSR-1 targets) to avoid silencing by piRNAs.

If self transcript licensing depends P granules, then these transcripts should be present in perinuclear condensates. Using RNA smFISH analyses, we examined the localization of *ceh-49*, an aberrantly silenced CSR-1 mRNA (Figure 6E), and noticed that *ceh-49* mRNAs were expressed in the pachytene region of the germline in wild type animals and both perinuclear and cytoplasmic signals are detected (Figure 6F). In *glh-1 glh-4* mutants, both perinuclear and cytoplasmic *ceh-49* signals were decreased. These results are consistent with the model that the P granule localization of some mRNA transcripts is critical for their protection from piRNA silencing (Ouyang et al. 2019).

We wondered why some CSR-1 targeted genes gained aberrant silencing in GLH/VASA mutants. We hypothesized that these genes might already be prone to silencing in wild type animals. To test this hypothesis, we compared the amounts of HRDE-1 associated 22G-RNAs to CSR-1 associated 22G-RNAs on these genes. Interestingly, in wild type animals, CSR-1 targets with enhanced HRDE-1 22G-RNA levels in *glh-1* mutants already exhibited a significantly higher ratio of HRDE-1 associated 22G-RNAs to CSR-1 associated 22G-RNAs compared to other CSR-1 targets (Figure 6G). This ratio for CSR-1 genes with enhanced HRDE-1 22G-RNA levels in *glh-1* mutants favors HRDE-1 even more in *glh-1* mutants (Figure 6G). These data indicate that these aberrantly silenced CSR-1 genes are already more targeted by silencing machinery than other CSR- 1 genes in wild type animals. In addition, previous studies have shown CSR-1 22G-RNAs are enriched at their 3’ end of transcripts (Ishidate et al. 2018; Singh et al. 2021). However, CSR-1 22G-RNAs are more distributed across the whole gene body for these aberrantly-silenced CSR-1 targets, similar to the distribution of CSR-1 22G-RNAs observed for WAGO targets (Figure S4C). Together, these observations suggest that perinuclear P granules are critical for protecting silencing-prone CSR-1 transcripts from aberrant piRNA silencing while at the same time promoting piRNA silencing on WAGO targets.

## DISCUSSION

Mutations in VASA helicase genes have been reported to lead to defects in the localization of PIWI in various animals (Malone et al. 2009)(Xiol et al. 2014)(Kuramochi-Miyagawa et al. 2010)(Elisabeth A. Marnik, J. Heath Fuqua, Catherine S. Sharp, Jesse D. Rochester, Emily L. Xu, Sarah E. Holbrook 2019)(Chen et al. 2020). Whether the localization of other small RNA factors in P granules and other perinuclear granules also relies on GLH/VASA helicases had only been explored sporadically. In *C. elegans*, piRNAs and other small RNA pathways factors are concentrated in distinct but partially overlapping granules, including P granules, Z granules and Mutator granules (Wan et al. 2018)(Phillips et al. 2012), suggesting these granules interact to facilitate gene regulation by small RNAs. Here we expand on these analyses to examine the roles GLH-1 and GLH-4 play in the localization of various small RNA factors, include P granule components - DEPS-1, EGO-1, CSR-1, Z granule components - WAGO-4 and ZNFX-1, and Mutator component MUT-16. We found that the formation of perinuclear condensation of these components are each compromised in *glh-1 glh-4* double mutants. Although not all the components examined are compromised to the same extent in *glh-1 glh-4* mutants, our observations suggest that GLH/VASA plays a global role in promoting the formation of these various liquid condensates.

Phase-separated condensates are capable of concentrating various proteins and RNAs, but whether these condensates indeed play a biological role remains controversial. For example, a previous study has reported that P body formation is a consequence but not a cause of miRNA-mediated gene silencing (Eulalio et al. 2007). In this study, we found that the production of piRNAs and other 22G-RNAs are not grossly affected in GLH/VASA mutants and other mutants defective in forming perinuclear condensates. Instead, the small RNA-based distinction of self and non-self RNAs are compromised in these mutants. Specifically, piRNA-dependent silencing of non-self is reduced, leading to increased mRNA expression. Simultaneously, hundreds of self RNAs (CSR-1 targets) were aberrantly silenced by piRNAs, leading to reduced mRNA expression. Together, these data argue that perinuclear P granules are critical for the fidelity of piRNA-based surveillance in *C. elegans* and provide an environment that allows small RNA factors to distinguish self from non-self. Our observations support a model where P granules act as the checkpoint for piRNA-mediated gene silencing, where the P granule promotes piRNA target recognition for non-self genes while allowing CSR-1 to guard self genes (Figure S4D). In GLH/VASA mutants, the P granule fails to form, leading to the dispersal of small RNA factors including PRG-1, WAGOs and CSR-1 into the cytoplasm. When denied the environment of the P granule, hundreds of typically silenced non-self transcripts fail to contact silencing machinery leading to their expression while hundreds of typically expressed self transcripts fail to contact CSR-1 leading to their repression by PRG-1. This model suggests P granules act as a specialized environment where transcripts are allowed residence time to properly contact either silencing or licensing machinery, as evidenced by our smFISH data that demonstrates the GLH/VASA-dependent perinuclear localization of both silenced and expressed mRNAs (Figure 2E and Figure 6F). Taken together, our study reveals the critical role of perinuclear P granules in promoting the fidelity of self and non-self nucleic acids distinction and has broad implications for the functions of other RNA-enriched liquid condensates.

## MATERIALS AND METHODS

### Caenorhabditis elegans strains

Animals were grown on standard nematode growth media (NGM) plates seeded with the Escherichia coli OP50 strain at 20°C or temperatures where indicated. Some strains were obtained from the Caenorhabditis Genetics Center (CGC).

The strains used in this study are glh-1(uoc1) I, glh-1^DQAD^(uoc3) I, glh- 1^FGGΔ^(uoc4) I, glh-1(uoc5) glh-4(gk225) I/hT2 [bli-4(e937) let-?(q782) qIs48] (I;III), deps-1(bn124) I, and meg-3(ax3055) meg-4(ax3052) X.

### Fluorescence microscopy and image processing

GFP- and RFP-tagged fluorescent proteins were visualized in living nematodes or dissected embryos by mounting young adult animals on 2% agarose pads with M9 buffer (22 mM KH2PO4, 42 mM Na2HPO4, and 86 mM NaCl) with 10-50 mM levamisole, or mounting one-cell embryos on 2% agarose pads by dissecting gravid hermaphrodites into egg salt buffer (5 mM HEPES pH 7.4, 118 mM NaCl, 40 mM KCl, 3.4 mM MgCl2, and 3.4 mM CaCl2). Fluorescent images were captured using a Zeiss LSM800 confocal microscope with a Plan-Apochromat 63X/1.4 Oil DIC M27 objective.

Image processing and quantification of fluorescent puncta were performed using ImageJ. Single slice images of gonads and images from maximum intensity projections of z-series of one-cell embryos were used for quantification. Regions of interest (ROIs) were selected, and areas of ROIs were measured. Image thresholds were set manually, and fluorescent puncta were selected. Puncta density was calculated as the number of fluorescent puncta in the ROI divided by the area of ROI. Integrated intensities and sizes of fluorescent puncta in all ROIs were measured. Images of 8-12 gonads were collected and quantified. Data were analyzed by student’s t-test or one-way ANOVA followed by Dunnett’s multiple comparisons.

### RNA isolation and quantitative real-time PCR

Total RNA was extracted using the standard method with TRIzol reagent (Invitrogen) from whole animals of ∼100,000 synchronized young adults. Stem-loop real-time PCR was performed to measure piRNA levels. 1 μg of total RNA was reverse transcribed with SuperScript IV Reverse Transcriptase (Invitrogen) in 1x reaction buffer, 5U SUPERase-In RNase Inhibitor (Invitrogen), 1 mM dNTPs, and 50 pM stem-loop reverse primer 5’- CTCAACTGGTGTCGTGGAGTCGGCAATTCAGTTGAG-n8-3’ (n8=reverse complement sequences of last 8 nucleotide acids in piRNAs). Each real-time PCR reaction consisted 4 μL of cDNA, 1 μM forward piRNA primer 5’- ACACTCCAGCTGGG-n16-3’ (n16= first 16 nucleotide acids in piRNAs), and 1 μM universal reverse primer 5’-CTCAAGTGTCGTGGAGTCGGCAA-3’. The amplification was performed using Power SYBR Green (Applied Biosystems) on the Bio-Rad CFX96 Touch Real-Time PCR Detection System. The experiments were repeated for a total of three biological replicates.

### Small RNA sequencing

Total RNA was extracted from whole animals of ∼100,000 synchronized young adults as described above. Small (<200nt) RNAs were enriched with mirVana miRNA Isolation Kit (Ambion). In brief, 80 μL (200-300 μg) of total RNA, 400 μl of mirVana lysis/binding buffer and 48 μL of mirVana homogenate buffer were mixed well and incubated at room temperature for 5 minutes. Then 176 μL of 100% ethanol was added and samples spun at 2500 x g for 4 minutes at room temperature to pellet large (>200nt) RNAs. The supernatant was transferred to a new tube and small (<200nt) RNAs were precipitated with pre-cooled isopropanol at -70°C. Small RNAs were pelleted at 20,000 x g at 4°C for 30 minutes, washed once with 70% pre-cooled ethanol, and dissolved with nuclease-free water. 10 μg of small RNA was fractionated on a 15% PAGE/7M urea gel, and RNA from 17 nt to 26 nt was excised from the gel. RNA was extracted by soaking the gel in 2 gel volumes of NaCl TE buffer (0.3 M NaCl, 10 mM Tris-HCl, 1 mM EDTA pH 7.5) overnight. The supernatant was collected through a gel filtration column. RNA was precipitated with isopropanol, washed once with 70% ethanol, and resuspended with 15 μL nuclease-free water. RNA samples were treated with RppH to convert 22G-RNA 5’ triphosphate to monophosphate in 1x reaction buffer, 10U RppH (New England Biolabs), and 20U SUPERase-In RNase Inhibitor (Invitrogen) for 3 hours at 37°C, followed by 5 minutes at 65°C to inactivate RppH. RNA was then concentrated with the RNA Clean and Concentrator-5 Kit (Zymo Research). Small RNA libraries were prepared according to the manufacturer’s protocol of the NEBNext Multiplex Small RNA Sample Prep Set for Illumina-Library Preparation (New England Biolabs). NEBNext Multiplex Oligos for Illumina Index Primers were used for library preparation (New England Biolabs). Libraries were sequenced using an Illumina HiSeq4000 to obtain single-end 36 nt sequences at the University of Chicago Genomic Facility.

### RNA immunoprecipitation sequencing (RIP-seq)

A total of ∼100,000 synchronized young adult animals were used for RIP-seq. Worm pellets were resuspended in equal volumes of immunoprecipitation buffer (20 mM Tris-HCl pH 7.5, 150 mM NaCl, 2.5 mM MgCl2, 0.5% NP-40, 80 U/mL RNase Inhibitor (Thermo Fisher Scientific), 1 mM dithiothreitol, and protease inhibitor cocktail without EDTA (Promega)), and grinded in a glass grinder for 8- 10 times. Lysates were clarified by spinning down at 15,000 rpm, 4°C, for 15 minutes. Supernatants were incubated with the GFP-Trap magnetic agarose beads (ChromoTek) at 4°C for 1 hour. Beads were washed with wash buffer (20 mM Tris-HCl pH 7.5, 150 mM NaCl, 2.5 mM MgCl2, 0.5% NP-40, and 1 mM dithiothreitol) six times, and then resuspended in TBS buffer for RNA extraction. Total RNA was extracted using the standard method with TRIzol reagent (Invitrogen). Small RNA libraries for RNA-seq were prepared as described above. Libraries were sequenced using an Illumina HiSeq4000 to obtain single-end 36 nt sequences at the University of Chicago Genomic Facility.

### Chemical cross-linking and co-immunoprecipitation of GLH-1

Chemical cross-linking of proteins was performed with Dithio-bismaleimidoethane (DTME). ∼100,000 synchronized flag::mCherry::GLH-1 young adults were collected and washed three times with M9. M9 was discarded to the same amount of the worm volume, then DTME dissolved in Dimethyl Sulfoxide (DMSO) was added to a final concentration of 2 mM. Samples were incubated for 30 minutes at room temperature with occasional shaking before washed three times with M9 to remove the cross-linker. Worm pellets were resuspended in equal volume of immunoprecipitation buffer (20 mM Tri-HCl pH 7.5, 150 mM NaCl, 2.5 mM MgCl2, 0.5% NP-40, 80 U/mL RNase Inhibitor (Thermo Fisher Scientific), 1 mM dithiothreitol, and protease inhibitor cocktail without EDTA (Promega)).

Worm pellets were homogenized using glass homogenizer for 15-20 strokes on ice. Lysates were centrifuged at 14000 x g (Eppendorf Centrifuge 5424 R) for 10 minutes to remove insoluble material. Supernatants were incubated with 25 μL of Anti-Flag Magnetic Beads (bioLinkedin) for 2 hours at 4°C on an end-to-end rotator. Supernatant was removed and beads were washed with 1 mL of wash buffer (20 mM Tris-HCl pH 7.5, 150 mM NaCl, 2.5 mM MgCl2, 0.5% NP-40) six times for 10 minutes each time, and with the final wash of 0.05% NP-40. Beads were incubated at 37°C for 30 minutes in 50 μL de-crosslinking buffer (50 mM Tris-HCl pH 7.5, 150 mM NaCl, 2 mM MgCl2, 0.2% Tween 20, 10 mM dithiothreitol). The final samples were boiled in 2 x SDS loading buffer at 100°C for 5 minutes before mass-spectrometry analysis.

### Mass-spectrometry analysis

The samples were subjected to SDS PAGE gel electrophoresis experiments, and gel slice was subjected to in-gel digestion with trypsin at overnight at 37°C. Purified peptides were desalted using a C18 Stage Tip column (Pierce), concentrated and dried. Then the peptides are reconstituted with 0.1% formic acid aqueous solution for subsequent liquid chromatography with tandem mass spectrometry (LC-MS/MS) analysis using a Q Exactive HF-X mass spectrometer (Thermo Scientific) equipped with a nanoliter flow rate Easy-nLC 1200 chromatography system (Thermo Scientific). The buffers used in the liquid chromatography separation consisted of buffer A (0.1% formic acid and 19.9% H2O) and buffer B (80% acetonitrile, 0.1% formic acid and 19.9% H2O). The liquid chromatography separation was achieved by linear gradient elution at a flow rate of 300 nL/min. The related liquid phase gradient was as follows: 0-3min, linear gradient of liquid B from 2% to 8%; 3-43 min, linear gradient of liquid B from 8% to 28%; 43-51 min, linear gradient of liquid B from 28% to 40%; 51-52 min, linear gradient of liquid B from 40% to 100%; 52-60 min, linear gradient of liquid B maintained at 100%. The Q Exactive HF-X mass spectrometer was used in the data dependent acquisition (DDA) mode. The full scan survey mass spectrometry analysis was performed as detection mode of positive ion, analysis time of 60 minutes, scan range of 350-1800 m/z, resolution of 60,000 at m/z 20, AGC target of 3e6, and maximum injection time (IT) of 50ms. The 20 highest intensity precursor ions were analyzed by the secondary mass spectrometry (MS2) with resolution of 15,000 at m/z 200, AGC target of 1e5, and maximum IT of 25ms, activation type of HCD, isolation window of 1.6 m/z, and normalized collision energy of 28. The mass spectrometry data were analyzed using MaxQuant 1.6.1.0 and used the database from Uniprot Protein Database (species: Caenorhabditis elegans).

### RNA smFISH

RNA smFISH was performed on dissected adult *C.elegans* gonads. For particular genotypes and conditions, experiments were performed with one or two technical replicates. Staged young adult worms were washed with M9 and resuspended in Dissection Buffer (PBS, 1mM EDTA) then deposited onto 18mm circular coverslips that were coated with 0.1% Poly-L-lysine solution. Gonads were dissected from whole worms using 25G x 5/8 hypodermic needles directly onto coverslips in a 12 well tissue culture plate. All subsequently steps were performed directly in the 12 well plate. Gonads were fixed in 4% formaldehyde for 30 minutes at room temperature. Gonads were then washed with PBS and dehydrated with 70% ethanol at 4C for one or two overnights. Gonads were re- hydrated with PBS and washed once with FISH wash buffer (SSC, 10% formamide) at 37C. FISH probes were suspended in Hybridization Buffer (10% formamide, 2mM vanadyl ribonucleoside complex, 20mg/mL BSA, 10mg/mL dextran sulfate, 2mg/mL *E.coli* tRNA) 1:50. smFISH probes were created by Biosearch Technologies to target *gfp* mRNA, *nos-3* mRNA, or *ceh-49* mRNA. *gfp* mRNA probes were conjugated to Quasar670, *nos-3* to CalFluor Red 610, and *ceh-49* to Quasar670. Gonads were hybridized overnight at 37C in 12 well tissue plate wrapped tightly in parafilm. Gonads were washed with FISH wash buffer, stained with DAPI, and mounted onto 25mm x 75mm x 1mm plain glass slides with ProLong Diamond Antifade Mountant. Samples were imaged on a Zeiss LSM800 Confocal Microscope at 40X magnification using a Plan-Apochromat 40X/1.4 Oil Objective.

### Sequencing data analysis

*RNA-seq*. Fastq reads were trimmed of adaptors using cutadapt (Martin 2011). Trimmed reads were aligned to the *C.elegans* genome build WS230 using bowtie2 ver 2.3.0 (Langmead and Salzberg 2012). After alignment, reads were overlapped with genomic features (protein-coding genes, pseudogenes, transposons) using bedtools intersect (Quinlan and Hall 2010). Reads per kilobase million (RPKM) values were then calculated for each individual feature by summing the total reads mapping to that feature, multiplied by 1e6 and divided by the product of the kilobase length of the feature and the total number of reads mapping to protein-coding genes. Protein-coding genes were used to normalize by sequencing depth because mRNA libraries were prepared by polyA tail selection, so reads mapping to features devoid of polyA tails are likely contaminants. RPKM values were then used in all downstream analyses using custom R scripts, which rely on packages ggplot2(Wickham 2016), reshape2 (Wickham 2007), ggpubr (Kassambara 2020), dplyr (Wickham et al. 2021). *sRNA-seq*. Fastq reads were trimmed using custom perl scripts. Trimmed reads were aligned to the *C.elegans* genome build WS230 using bowtie ver 1.2.1.1 (Langmead et al. 2009) with options -v 0 --best --strata. After alignment, reads that were between 17-40 nucleotides in length were overlapped with genomic features (rRNAs, tRNAs, snoRNAs, miRNAs, piRNAs, protein-coding genes, pseudogenes, transposons) using bedtools intersect (Quinlan and Hall 2010). Sense and antisense reads mapping to individual miRNAs, piRNAs, protein-coding genes, pseudogenes, RNA/DNA transposons, simple repeats, and satellites were totaled and normalized to reads per million (RPM) by multiplying be 1e6 and dividing read counts by total mapped reads, minus reads mapping to structural RNAs (rRNAs, tRNAs, snoRNAs) because these sense reads likely represent degraded products. Reads mapping to multiple loci were penalized by dividing the read count by the number of loci they perfectly aligned to. Reads mapping to miRNAs and piRNAs were only considered if they matched to the sense annotation without any overlap. In other words, piRNA and miRNA reads that contained overhangs were not considered as mature piRNAs or miRNAs respectively. 22G-RNAs were defined as 21 to 23 nucleotide long reads with a 5’G that aligned to protein-coding genes, pseudogenes, or transposons. RPM values were then used in all downstream analyses using custom R scripts using R version 4.0.0 (R Core Team 2020), which rely on packages ggplot2 (Wickham 2016), reshape2 (Wickham 2007), ggpubr (Kassambara 2020), dplyr (Wickham et al. 2021). To determine GLH-dependent 22G-RNA lists, a Bayesian approach as in (Maniar and Fire 2011).

*Metagene analysis.* Metagene profiles were calculated by computing the depth at each genomic position using 21 to 23 nucleotide long small RNA reads with a 5’G using bedtools genomecov (Quinlan and Hall 2010). A custom R script was then used to divide genes into 100 bins and sum the normalized depth within each bin. Groups of genes were then plotted using the sum of the normalize depth at each bin.

## ACKNOWLEDGEMENTS

We thank Dr. Edwin Ferguson, Dr. Karen Bennett, and members of the Lee lab for critical comments on the manuscript. Some strains used in this study were provided by the CGC, which is funded by NIH Office of Research Infrastructure Programs (P40 OD010440). This work is supported in part by NIH predoctoral training grant T32 GM07197 to J.B.; the National Center for Advancing Translational Sciences (NCATS) of the National Institutes of Health (NIH) grant 1UL1TR002389-01 to the Institute for Translational Medicine (ITM); the National Natural Science Foundation of China (grants 31771500 and 31922019) and the program for HUST Academic Frontier Youth Team (grant 2018QYTD11) to DZ; the Ministry of Science of Technology of Taiwan (MOST 108-2628-E-006-004- MY3 and MOST 110-2221-E-006-198-MY3 grants to W.-S.W. the NIH P01 grant (HD078253) to Z.W, the NIH grant R01-GM132457 to H.-C.L.

---

**Figure S1.**
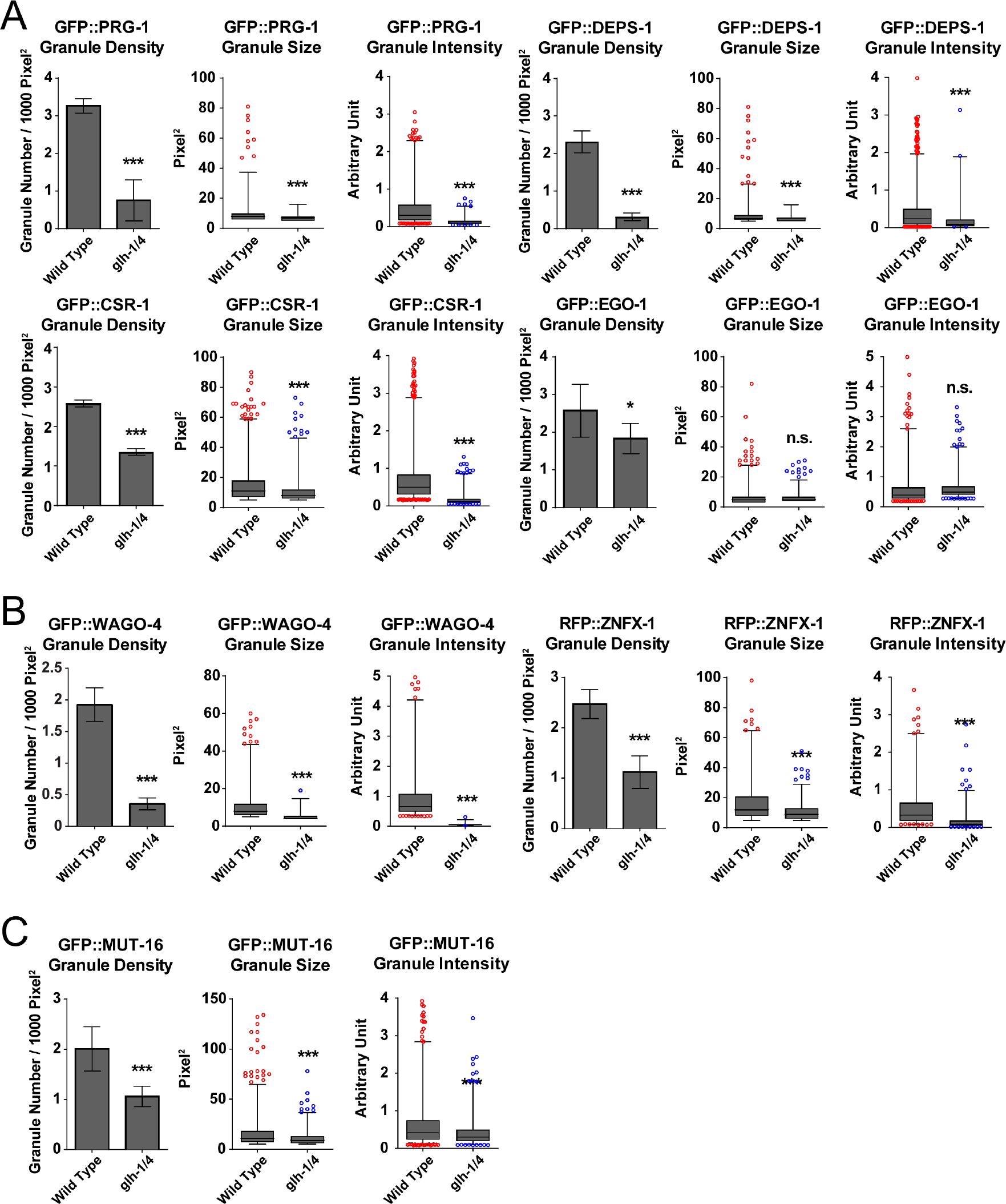
Image analyses corresponding to Figure 1A-C. (a) Granule density, size, and intensity quantification of proteins known to be enriched in P granules in the wild type or the *glh-1 glh-4* mutant, corresponding to micrographs in Figure 1B. Each bar represents n = 8∼10 worms. (b) Granule density, size, and intensity quantification of proteins known to be enriched in Z granules in the wild type or the *glh-1 glh-4* mutant, corresponding to micrographs in Figure 1C. Each bar represents n = 8∼10 worms. (c) Granule density, size, and intensity quantification of proteins known to be enriched in mutator granules in the wild type or the *glh-1 glh-4* mutant, corresponding to micrographs in Figure 1D. Each bar represents n = 8∼10 worms. For a-c, statistical analysis was performed using a one-tailed Student’s t-test. *** indicates *P* < 0.001.

**Figure S2.**
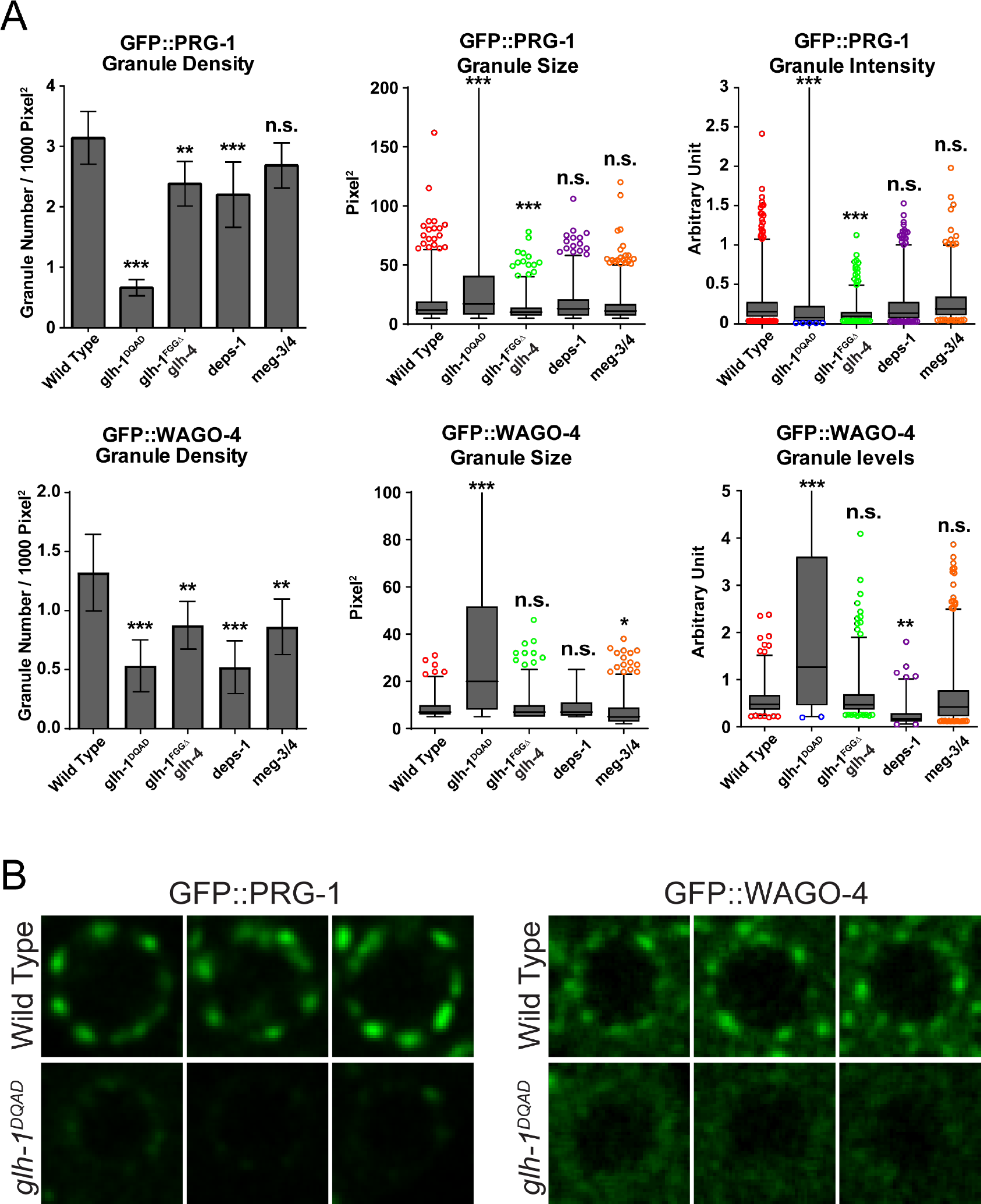
Image analyses corresponding to Figure 2B. (a) Granule density, size, and intensity quantification of GFP::PRG-1 and GFP::WAGO-4 in the wild type or the *glh-1 glh-4* mutant, corresponding to micrographs in Figure 1B. Each bar represents n = 8∼10 worms. statistical analysis was performed using a one-tailed Student’s t-test. *** indicates *P* < 0.001. (b) Single nuclei from micrographs presented in Figure 2B.

**Figure S3.**
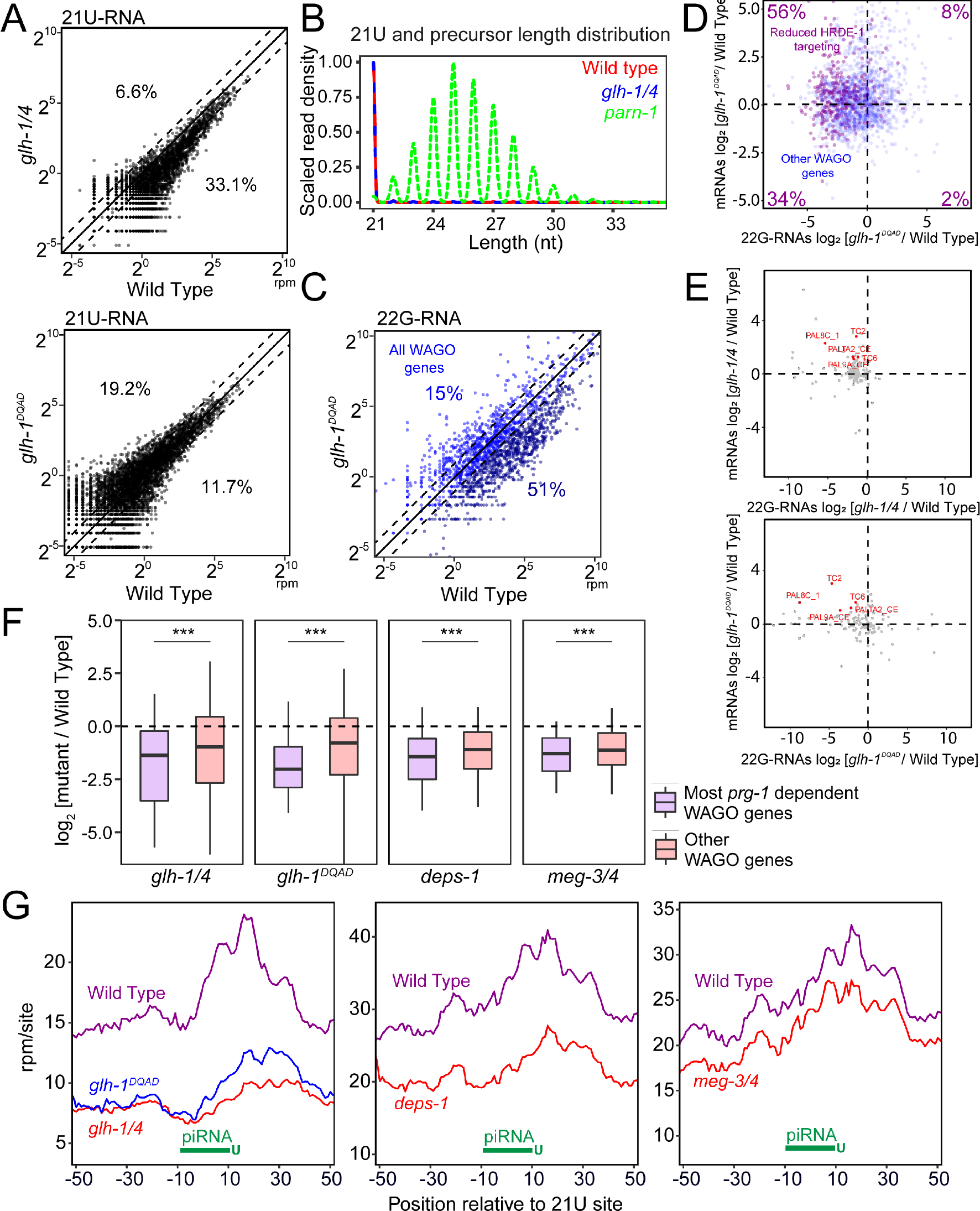
GLH/VASA mutants exhibit defects in producing secondary WAGO-22G-RNAs (Corresponding to Figure 4) (A) A scatter plot showing the abundance of each piRNA (21U-RNA) in *glh-1 glh-4* and *glh-1* DQAD mutant worms compared to wild type. The three diagonal lines indicate a two-fold increase (top), no change (middle), or a two-fold depletion (bottom) in the indicated mutant strains. Percentages indicate the proportion of piRNAs increased or decreased 2-fold compared to wild type. (B) Scaled kernel density estimation of mature piRNAs (21nt) and their precursors (>21nt) in the *glh-1 glh-4* double mutant, the *parn-1* mutant and wild type animals. (C) A scatter plot showing the abundance of all 22G-RNAs mapped to each WAGO target in wild type worms compared to *glh-1 DQAD* mutants. The percentage of WAGO targets with 2-fold increased or decreased 22G- RNAs in mutants are shown. The three diagonal lines indicate a two-fold increase (top), no change (middle), or a two-fold depletion (bottom) in *glh- 1 DQAD* mutants. (D) A scatter plot showing the mRNA vs 22G-RNA log2 expression changes for WAGO targets in *glh-1 DQAD* mutants vs wild type worms. The upper left quadrant corresponds to WAGO targets that have become activated in the mutant (increased mRNAs and decreased 22G-RNAs). Percentages are shown in each quadrant to indicate the proportion of WAGO targets with reduced HRDE-1-associated 22G-RNAs in the *glh-1* mutant that fall in that quadrant. (E) A scatter plot showing the mRNA vs 22G-RNA log2 expression changes for transposon mRNA in the indicated mutants vs wild type worms. The upper left quadrant corresponds to targets that have become activated in the mutant (increased mRNAs and decreased 22G-RNAs). Transposons with increased mRNA and decreased 22G-RNAs in both *glh-1 glh-4* and *glh-1 DQAD* mutants are highlighted in red. (F) 22G-RNA fold changes for WAGO targets most dependent on *prg-1* for 22G-RNA accumulation versus all other WAGO targets in the indicated mutants compared to wild type. Most *prg-1* dependent WAGO targets defined as those WAGO targets with a significant (*P* < 0.05 and 2-fold) decrease in 22G-RNA accumulation in *prg-1* mutants. Statistical analysis was performed using a two-tailed Mann-Whitney Wilcoxon test. *** indicates *P* < 0.001. (G) Density of 22G-RNAs within a 100 nt window around predicted piRNA target sites in the indicated strains. Computed by summing 22G-RNA density per piRNA targeting site in all WAGO targets. The plots are centered on the 10^th^ nucleotide of piRNAs.

**Figure S4.**
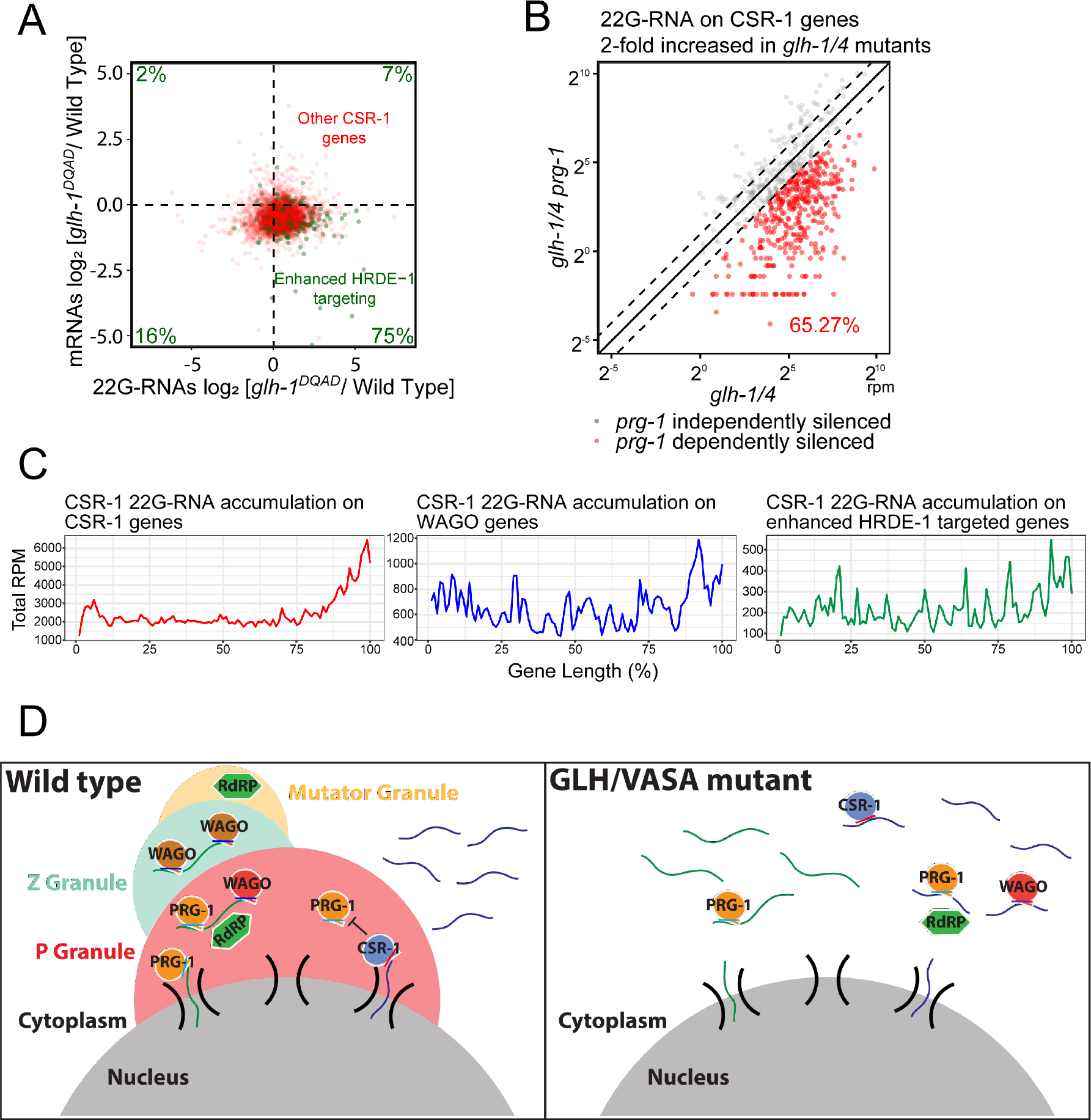
Many functional germline genes are silenced in mutants with defects in forming perinuclear PRG-1 granules (Corresponding to Figure 6) (A) A scatter plot showing the mRNA vs 22G-RNA log2 expression changes for CSR-1 targets in *glh-1 DQAD* mutants vs wild type worms. The bottom right quadrant corresponds to CSR-1 targets that have become silenced in the mutant (decreased mRNAs and increased 22G-RNAs). Percentages are shown in each quadrant to indicate the proportion of CSR-1 targets with enhanced HRDE-1-associated 22G-RNAs in the *glh-1* mutant that fall in that quadrant. (B) A scatter plot showing 22G-RNA accumulation on CSR-1 genes with a 2- fold increase in 22G-RNA accumulation in *glh-1 glh-4* mutants. The *glh-1 glh-4 prg-1* triple mutant is compared to the *glh-1 glh-4* double mutant. In red, enhanced CSR-1 targets that are suppressed by *prg-1* loss are shown. The three diagonal lines indicate a two-fold increase (top), no change (middle), or a two-fold depletion (bottom) in the indicated mutant strains. (C) Metagene traces showing the total accumulation of 22G-RNAs by percentage of gene length in CSR-1 IP experiments. Traces are shown for three classes of genes: all CSR-1 targets (left), all WAGO targets (middle), and CSR-1 targets with enhanced HRDE-1 associated 22G- RNAs in the *glh-1* mutant (right). (D) Model showing the P granule working in concert with Z and Mutator granules to maintain an environment that protects specific self genes from silencing and promotes the silencing of specific non-self genes (Left). In GLH/VASA mutants where granule integrity is lost, hundreds of typically expressed transcripts become targeted by PRG-1 and silencing machinery while hundreds of typically silenced transcripts become dispersed into the cytoplasm and expressed (Right).

